# Translational repression of the *Drosophila nanos* mRNA involves the RNA helicase Belle and RNA coating by Me31B and Trailer hitch

**DOI:** 10.1101/141655

**Authors:** Michael Götze, Jérémy Dufourt, Christian Ihling, Christiane Rammelt, Stéphanie Pierson, Nagraj Sambrani, Claudia Temme, Andrea Sinz, Martine Simonelig, Elmar Wahle

**Author notes:** current address: IDT, Am Pharmapark, 06861 Dessau-Rosslau, Germany.

## Abstract

Translational repression of maternal mRNAs is an essential regulatory mechanism during early embryonic development. Repression of the *Drosophila nanos* mRNA, required for the formation of the anterior-posterior body axis, depends on the protein Smaug binding to two Smaug recognition elements (SREs) in the *nanos* 3’ UTR. In a comprehensive mass-spectrometric analysis of the SRE-dependent repressor complex, we identified Smaug, Cup, Me31B, Trailer hitch, eIF4E and PABPC, in agreement with earlier data. As a novel component, the RNA-dependent ATPase Belle (DDX3) was found, and its involvement in deadenylation and repression of *nanos* was confirmed *in vivo*. Smaug, Cup and Belle bound stoichiometrically to the SREs, independently of RNA length. Binding of Me31B and Tral was also SRE-dependent, but their amounts were proportional to the length of the RNA and equimolar to each other. We suggest that ‘coating’ of the RNA by a Me31B•Tral complex may be at the core of repression.

## Introduction

Control of gene expression by translational regulation of mRNAs is found throughout biology, but particularly important in oocyte development and early embryogenesis in animals. As the zygotic genome is not transcribed during very early development, mRNAs required at this time are produced during oocyte development (maternal mRNAs), and many are stockpiled in a repressed, ‘masked’ state. During maturation of the oocyte to a fertilizable egg and the first phases of embryonic development, specific maternal mRNAs are translationally activated in a controlled manner. Many are also regulated by localization at specific sites and by degradation (Lasko 2011; Barckmann and Simonelig 2013; Laver et al. 2015).

In *Drosophila*, zygotic genome activation is a gradual process; full-scale zygotic transcription does not commence until nuclear division cycle 14, with the beginning of the cellular blastoderm stage (Ali-Murthy et al. 2013; Harrison and Eisen 2015; Laver et al. 2015). One maternal RNA governing early development is the *nanos (nos)* mRNA. Its regulation is essential for development: Formation of the anterior-posterior axis of the embryo depends on the Nos protein being produced exclusively at the posterior pole (Wang and Lehmann 1991). For this purpose, most of the *nos* mRNA, which is distributed throughout the embryo, is translationally repressed (Gavis and Lehmann 1994) and degraded over the first 2–3 h of development (Dahanukar and Wharton 1996; Bashirullah et al. 1999). At most ~4% of the *nos* mRNA is localized at the posterior pole (Bergsten and Gavis 1999; Trcek et al. 2015) and, due to stabilization and derepression by Oskar, serves as a localized source of Nos (Ephrussi and Lehmann 1992; Smith et al. 1992; Dahanukar et al. 1999). Both repression and degradation of non-localized *nos* mRNA depend on the protein Smaug (Smg) (Dahanukar and Wharton 1996; Smibert et al. 1996; Dahanukar et al. 1999; Smibert et al. 1999) and the Piwi-interacting RNA (piRNA) machinery (Rouget et al. 2010). Smg is essential for the maternal-to-zygotic transition (Benoit et al. 2009), causing repression and degradation of hundreds of maternal mRNAs (Tadros et al. 2007; Chen et al. 2014a). Smg regulates *nos* by binding two Smaug Recognition Elements (SREs) in the *nos* 3’ UTR and recruits the CCR4-NOT complex, which catalyzes mRNA deadenylation (Semotok et al. 2005; Jeske et al. 2006; Zaessinger et al. 2006). For translational repression, Smg binds the protein Cup (Nelson et al. 2004), and a miRNA-independent repressive role of Ago1 has also been reported (Pinder and Smibert 2013), but the mechanism of repression is not fully understood (Jeske et al. 2011).

Deadenylation and translational repression of *nos* can be observed in extracts from early *Drosophila* embryos (Jeske et al. 2006; Jeske et al. 2011). Deadenylation and repression both depend on the SREs and, by inference, on Smg, but are independent of each other. Smg-associated Cup inhibits translation by binding the cap-binding translation initiation factor eIF4E and competitively displacing eIF4G (Nelson et al. 2004; Jeske et al. 2011). However, the 5’ cap as well as eIF4E and eIF4G are dispensable for SRE-dependent repression (Jeske et al. 2011); thus, an additional repression mechanism must exist. In support of this, the SRE-dependent repressor complex contains the proteins Me31B and Trailer hitch (Tral) in addition to Smg, Cup and eIF4E (Jeske et al. 2011). Me31B and its orthologues are DEAD box family RNA helicases/RNA-dependent ATPases and involved in translational repression in flies (Nakamura et al. 2001; Tritschler et al. 2009), yeast (Dhh1p) (Coller and Parker 2005) and vertebrates (DDX6/p54/RCK) (Minshall et al. 2001; Chen et al. 2014b; Mathys et al. 2014). Tral *(S. cerevisiae* Scd6p; *C. elegans* CAR1; vertebrate Rap55 or Lsm14) associates with Me31B and also represses translation (Audhya et al. 2005; Boag et al. 2005; Wilhelm et al. 2005; Tanaka et al. 2006; Weston and Sommerville 2006; Nissan et al. 2010; Hubstenberger et al. 2013; Ayache et al. 2015).

Formation of the SRE-dependent repressor complex in embryo extract is ATP-dependent and slow, requiring 20–30 minutes. Once formed, the complex is kinetically unusually stable, with an estimated t_1/2_ of ~ 4 h. These observations suggest that the repressor complex is not governed by a simple association-dissociation equilibrium. Presumably due to the stability of the complex, repression is ~ twenty-to fiftyfold, i. e. ≥ 95% of the SRE-containing RNA is turned off. Importantly, SRE-dependent repression acts on translation initiation driven by the CRPV IRES. As this IRES can directly associate with ribosomes, independently of any initiation factors, the repressor complex likely affects either ribosome association or elongation (Jeske et al. 2011).

Here we report a systematic analysis of the composition of the SRE-dependent repressor complex. In addition to the previously known proteins, the DEAD-box protein Belle (Bel) was found in the complex. Genetic experiments confirmed that Bel participates in *nos* regulation *in vivo*. Me31B and Tral bind in multiple copies along the repressed RNA, presumably sequestering it in a form that is inaccessible for ribosomes.

## Results

### Composition of the SRE-dependent repressor complex

For an analysis of the constituents of the SRE-dependent repressor complex, gradient centrifugation was used as a first purification step. In **Fig. 1**, radiolabeled, m7G-capped luciferase RNAs were used carrying a *nos* 3’ UTR fragment with two SREs, either wild type (SRE^+^) or with an inactivating point mutation in each (SRE^+^). The RNAs had ‘internal’ poly(A) tails, which stimulate translation like a 3’-terminal tail, but are protected from SRE-dependent deadenylation by flanking 3’ sequences (Jeske et al. 2011) (**Fig. 1A**). The RNAs were incubated in *Drosophila* embryo extract under conditions allowing formation of the repressor complex (first preincubation) and then sedimented through a sucrose gradient (**Fig. 1B, C**). Aliquots of the peak fractions were assayed for translation in embryo extract either directly or after a second preincubation with fresh extract. RNA present in the gradient fractions was strongly repressed without the second preincubation. As a control, untreated luciferase RNA required a preincubation in order to develop the full extent of repression (**Fig. 1B, D**). Thus, the repressor complex formed during the first preincubation survived the long fractionation procedure, in agreement with its known kinetic stability (Jeske et al. 2011). The RNA sedimented faster than in earlier experiments showing SRE-dependent inhibition of 48S complex formation (Jeske et al. 2011), probably due to differences in experimental conditions. With precautions taken to suppress RNase activity in the extract (see Materials and Methods), RNA stability was not significantly different between the repressed SRE^+^ RNA and the SRE^−^ control (**Fig. 1E, F**).

**Fig. 1:**
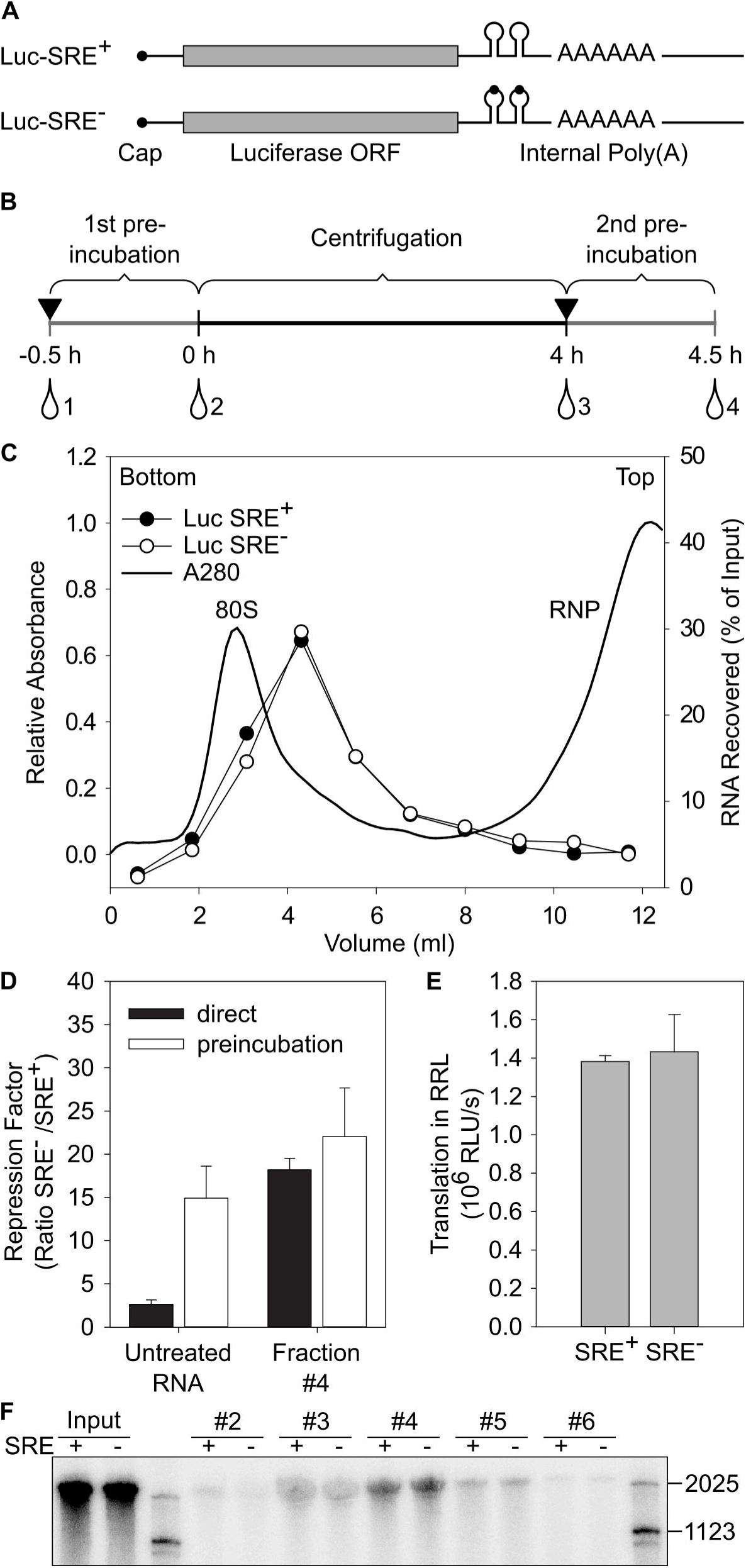
Reporter RNAs maintain their repressed state during gradient centrifugation. (A) Cartoon of luciferase reporter RNAs. (B) Scheme of the assay. Black triangels indicate addition of embryo extract, and drop symbols indicate samples withdrawn for translation and luciferase assays. Numbers refer to the data shown in (**D**). (C) Repressor complexes formed on radiolabelled reporter RNAs were separated by sucrose gradient sedimentation. Distributions of the two RNAs are overlayed. UV absorption indicates the positions of free RNPs and the 80S ribosome. (D) As shown in (B), luciferase RNAs were tested for translational repression either directly or after preincubation in embryo extract (‘untreated’; samples 1 and 2 in [B]). A second set of samples was preincubated in extract and then separated by gradient centrifugation. Aliquots from the peak fractions as in (**C**) were assayed for translation in embryo extract either with or without a second preincubation in fresh extract (‘fraction #4’; samples 3 and 4 in [**B**]). (E) RNAs were purified from equal volumes of the peak fractions of gradients as in (**C**), and equal aliquots were assayed for translation in rabbit reticulocyte lysate, which does not exhibit SRE-dependent repression (Jeske et al. 2006). Thus, similar luciferase yields indicated similar RNA recoveries for both RNAs. Error bars represent the standard deviation of three independent experiments. (F) Radiolabelled luciferase RNA from the sucrose gradient shown in **Fig. 1C** was purified and analysed by denaturing gel electrophoresis and phosphorimaging. Numbers above the lanes indicate fraction numbers of the sucrose gradient. Note that the inclusion of ‘short RNA’ (see Materials and Methods) strongly stabilized the RNA compared to earlier experiments (Jeske et al. 2011).

For the actual analysis of the repressor complex, similar RNAs as in **Fig. 1** were used (1-AUG *nos* and 1-AUG *nos* SRE^−^) that only differed by containing a shorter ORF and no 5’ cap; the cap is irrelevant for repression and stability of the repressor complex (Jeske et al. 2011). The RNAs were randomly biotinylated. After incubation in extract, these shorter RNAs showed an SRE-dependent difference in sedimentation, the SRE^+^ RNA sedimenting in the 80S region ahead of the control; this was more visible in an analytical gradient (**Fig. 2A**) than in the preparative experiment (**Fig. 2B**). Gradient fractions were selected as shown in **Fig. 2B**, pooled from several runs and concentrated. Equal quantities of the SRE^+^ and SRE^-^ RNPs, based on trace-labeling of the RNA, were affinity-purified on streptavidin beads, and proteins eluted by SDS were analyzed by gel electrophoresis (**Fig. 2C**) followed by liquid chromatography/tandem mass spectrometry (LC/MS/MS). Proteins detected were evaluated by label-free quantification (MaxQuant) (Cox and Mann 2008) based on the intensities of the MS signals and spectral counts, corrected for the molecular mass of each protein. In **Fig. 2E**, the apparent abundance of each protein is plotted for the SRE^+^ RNA against the control. All proteins are listed in **Supplemental Table 1**.

**Fig. 2:**
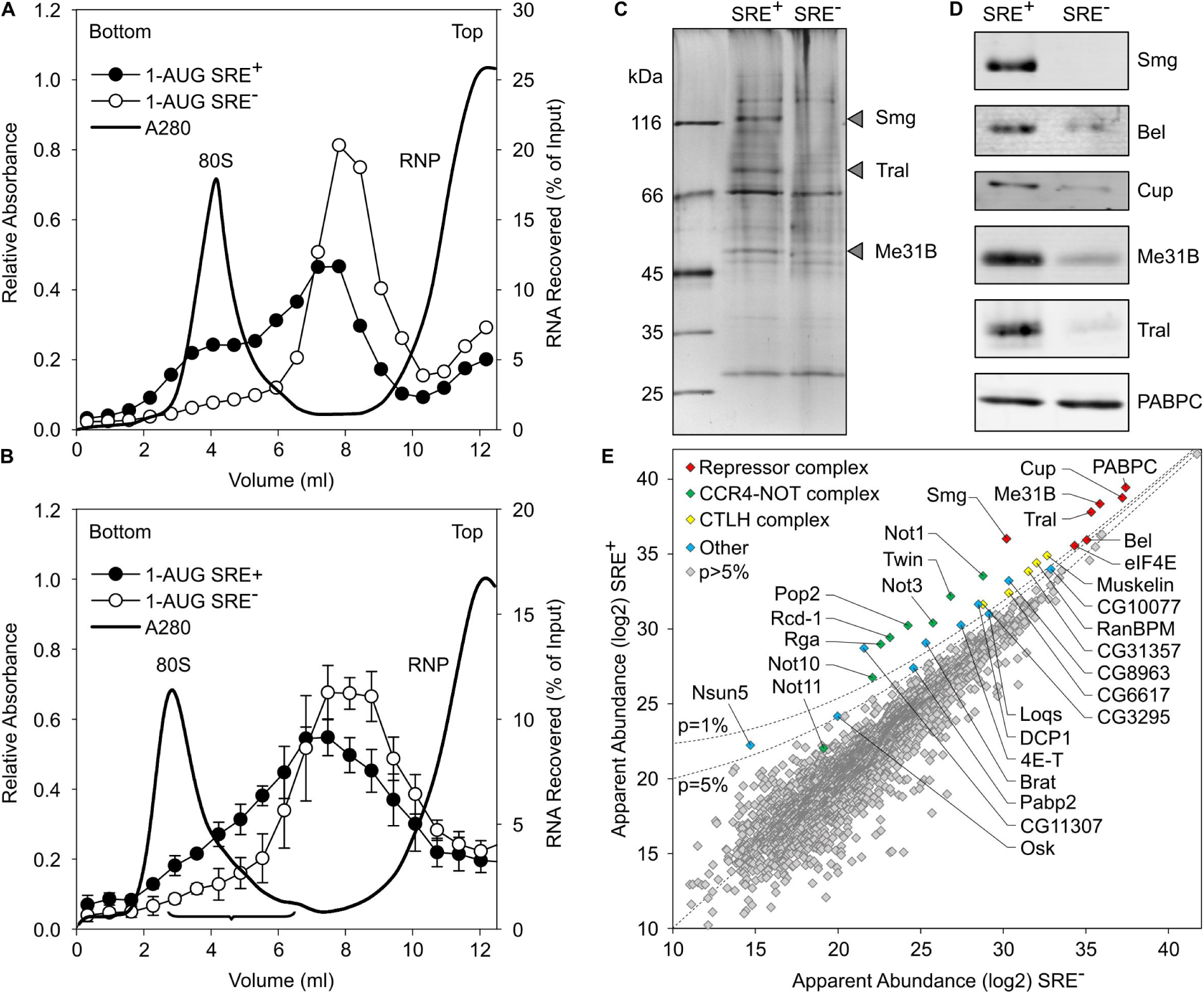
Analysis of the SRE-dependent repressor complex (A) Radiolabelled, biotinylated RNAs (1-AUG *nos* and 1-AUG *nos* SRE^−^) were incubated for assembly of a repressor complex and separated on a sucrose gradient as in Fig. (B), but the volume loaded was smaller (0.2 ml versus 1 ml). (B) Radiolabelled, biotinylated RNAs (1-AUG *nos* and 1-AUG *nos* SRE^−^) were separated on a preparative sucrose gradient (see Materials and Methods). Fractions pooled for the analysis of the repressor complex are indicated by the bracket. Error bars represent the standard deviation (n=4). (C) Corresponding fractions from a total of 12 gradients from 4 independent experiments each for the SRE^+^ RNA and the SRE^−^ control were pooled, and RNPs were purified on streptavidin beads. Equal amounts based on trace-labeling of the RNA were analysed by SDS-PAGE and silver-staining. Arrow heads indicate bands enriched in the SRE^+^ RNP that might correspond to Smaug (109 kDa), Trailer Hitch (69 kDa) and Me31B (52 kDa). (D) Specific association of proteins with the SRE^+^ RNA was confirmed by western analysis in an independent pull-down assay. Smg, Me31B and PABPC served as controls for Bel. (E) Proteins in the purified RNP fractions were analyzed by mass spectrometry and label-free quantification. Apparent protein abundance in SRE^+^ versus SRE^−^ RNP was plotted on a log2 scale. Proteins enriched in the SRE^+^ RNP beyond p = 0.05 and NOT11 are labeled. The complete list of proteins represented in (E) is found in **Table S1**. Different sets of proteins in the same data are highlighted in **Fig. S1**, and additional enriched proteins are listed in **Table S2**.

As expected, Smg was among the most abundant proteins and most strongly enriched in the SRE^+^ RNP. Cup, Tral and Me31B formed a tight cluster with an apparent abundance even higher than Smg and enriched in the SRE^+^ RNP. Three additional proteins were also abundant and enriched in the SRE^+^ RNP: First, enrichment of eIF4E-1 (Hernandez et al. 2005) agrees with previous results (Jeske et al. 2011). As the RNA was not capped, the protein’s presence was presumably due to protein-protein interactions, e. g. with Cup (Nelson et al. 2004; Chekulaeva et al. 2006) and Me31B (Minshall and Standart 2004). Second, the presence of PABPC was expected due to the internal poly(A) tail. An SRE-dependent enrichment of the protein agrees with the observation that a poly(A) tail facilitates repression (Jeske et al. 2006; Jeske et al. 2011) and with the presence of PABPC in DDX6 complexes purified under stringent conditions (Ayache et al. 2015; Bish et al. 2015). In contrast, western analyses of RNP complexes isolated by a simple pull-down procedure consistently showed the PABPC content to be independent of the SREs (Jeske et al. 2011) (**Fig. 2D**). The procedure leading to the MS analysis took considerably longer than a simple pull-down and might thus reveal a more stable association of PABPC with the repressed RNA compared to the control. Third, a novel component, the RNA-dependent ATPase Belle (Bel) (Johnstone et al. 2005) was identified. Its specific association with the SRE^+^ RNA was confirmed by western blot (**Fig. 2D**, **Fig. 5**).

The CCR4-NOT complex is responsible for Smg-dependent deadenylation (Semotok et al. 2005; Zaessinger et al. 2006) and associates with SRE-containing RNAs (Jeske et al. 2011). Satisfyingly, all core components of the complex (Not1, Ccr4/Twin, Caf1/Pop2, Not2/Rga, Not3 and Caf40/Rcd-1) formed a cluster of similar SRE-specific enrichment and roughly similar abundance; enrichment of NOT10 was less pronounced, and Not11 was even weaker (**Fig. 2E**). However, reduced abundance of all subunits compared to the Smg/Cup cluster suggests that the CCR4-NOT complex is not part of the stable core of the repressor complex. Several other proteins were enriched in the SRE^+^ RNP, but less so than either the Smg/Cup cluster or the CCR4-NOT complex (**Fig. 2E**; **Table S2**). A low-level presence of Oskar may be related to its role in derepression of *nos* in the pole plasm. The relationship of the other modestly abundant and enriched proteins to the repressor complex is uncertain. Among them, five subunits of the conserved CTLH (C-terminal to LisH [Lissencephaly type-1-like homology motif]) complex (Francis et al. 2013) stood out.

As expected (Nelson et al. 2004; Jeske et al. 2011), eIF4G was moderately depleted from the repressed RNP. All other initiation factors were less abundant than eIF4E and eIF4G and not enriched in either RNP (**Fig. S1A**). Ribosomal proteins were depleted from the SRE^+^ RNP (**Fig. S1B, D**). Ago 1 has been reported to participate in SRE-dependent repression (Pinder and Smibert 2013), but was not enriched in the SRE^+^ RNP. Ago2, the GW182 protein Gawky, or proteins involved in the piRNA pathway (Piwi, Aubergine, Ago3) were not enriched either (**Fig. S1C**). This is not unexpected as the region of the *nos* 3’ UTR most strongly targeted by piRNAs (Rouget et al. 2010; Barckmann et al. 2015) was not present in our constructs.

Pat1 (HPat or Patr-1 in *Drosophila)* and EDC3 bind the same surface of Me31B as Tral (Tritschler et al. 2008; Tritschler et al. 2009; Haas et al. 2010; Jonas and Izaurralde 2013; Sharif et al. 2013). Consistent with this competition, EDC3 and HPat were present at much lower levels than Tral and weakly enriched on the SRE^+^ RNA (**Fig. S1C**). A Me31B-Tral-Cup complex also contains the decapping complex component Dcp1, but not the catalytic subunit, Dcp2 (Tritschler et al. 2008). Dcp1 was enriched in the SRE-dependent RNP, but not abundant. Dcp2 was not detected at all. Ypsilon schachtel (Yps) and Exuperantia (Exu) have been found in Me31B-containing RNPs (Nakamura et al. 2001; Wilhelm et al. 2005), but Yps was not enriched in the repressor complex (**Fig. S1C**). Western blotting confirmed an equal association with either RNA (data not shown). Exu was not detected, consistent with the absence of *nos* from immunoprecipitated Exu-Yps complexes (Wilhelm et al. 2000).

In an independent experiment, Smg was immunoprecipitated from extract that had not been treated with RNase. Proteins were identified by LC/MS/MS and compared to a pre-immune serum control. The results supported those of the streptavidin purification: Core components of the repressor complex were enriched with Smg; only the enrichment of Belle was weak. All core subunits of the CCR4-NOT complex and four subunits of the CTLH complex were also enriched (**Fig. 3**, **Table S3**).

**Fig. 3:**
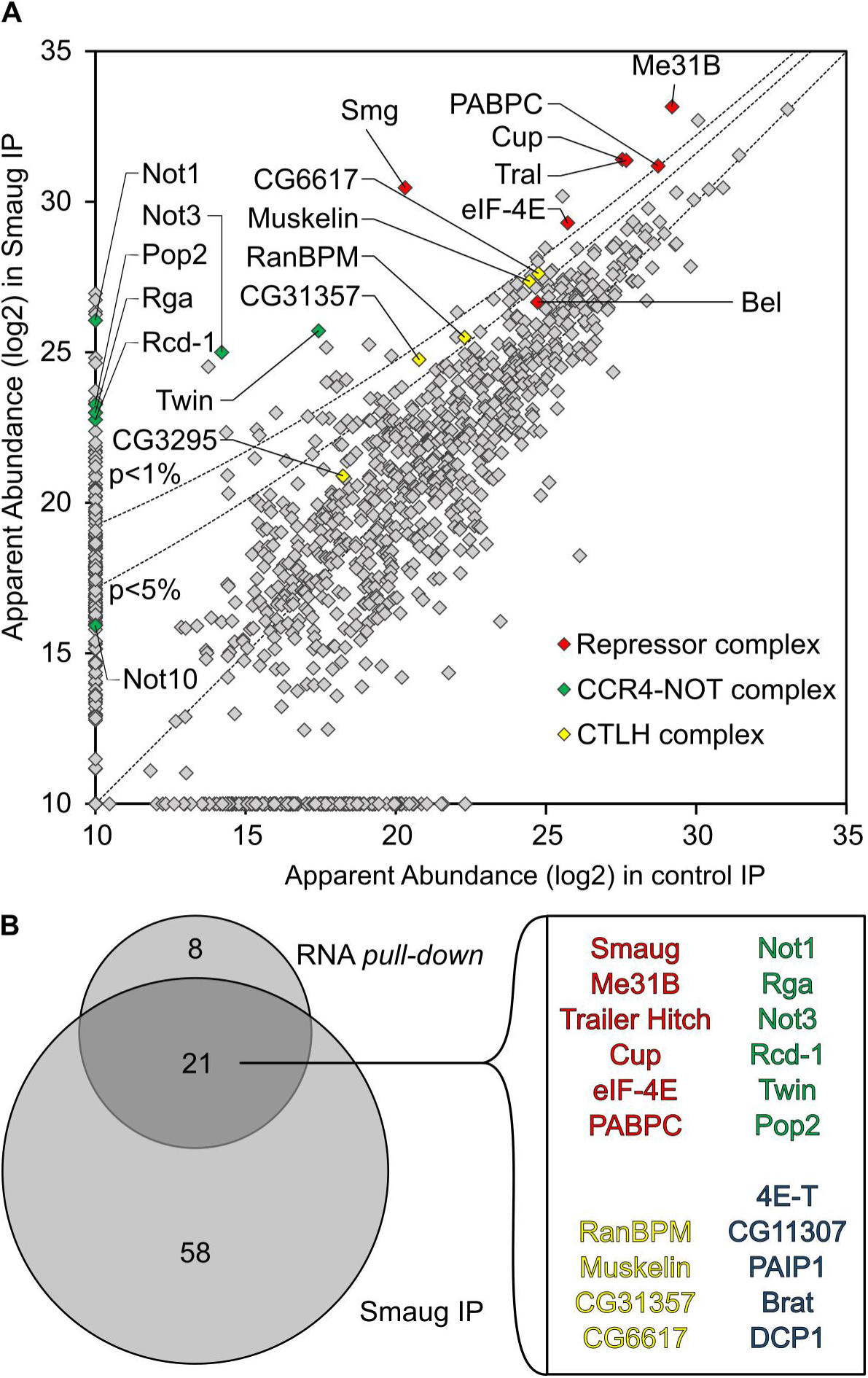
MS analysis of proteins co-precipitated with Smg. (A) Quantitative MS data were plotted for the Smg immunoprecipitation versus a preimmune control. Proteins that were also significantly enriched in the strepavidin pull-down of the SRE-dependent repressor complex are highlighted as in **Fig 2E**. p value cutoffs are indicated as lines. (B) Venn diagram comparing proteins enriched beyond p = 0.05 in the Smg immunoprecipitation and in the streptavidin pull-down (**Fig. 2E**). The 21 proteins in the overlap are listed. Belle, NOT10 and the CTLH complex subunit CG3295 had p values higher than 0.05. All proteins enriched in the Smg IP are listed in **Table S3**.

### Repressed *nos* mRNA exists as a monomeric RNP

The repressed RNPs sedimented rapidly, comparable to ribosomes (**Fig. 2A**). In the case of *oskar* mRNA, oligomerization of the repressed RNPs contributes to their rapid sedimentation (Chekulaeva et al. 2006; Besse et al. 2009). However, when a biotinylated SRE^+^ RNA was incubated in embryo extract together with a second SRE^+^ RNA, lacking biotin and distinguishable by size, streptavidin pull-down resulted in the purification of only the biotinylated RNA; no association with the second RNA was seen (**Fig. 4A**). We conclude that Smg-dependent repression does not involve RNA oligomerization.

**Fig. 4:**
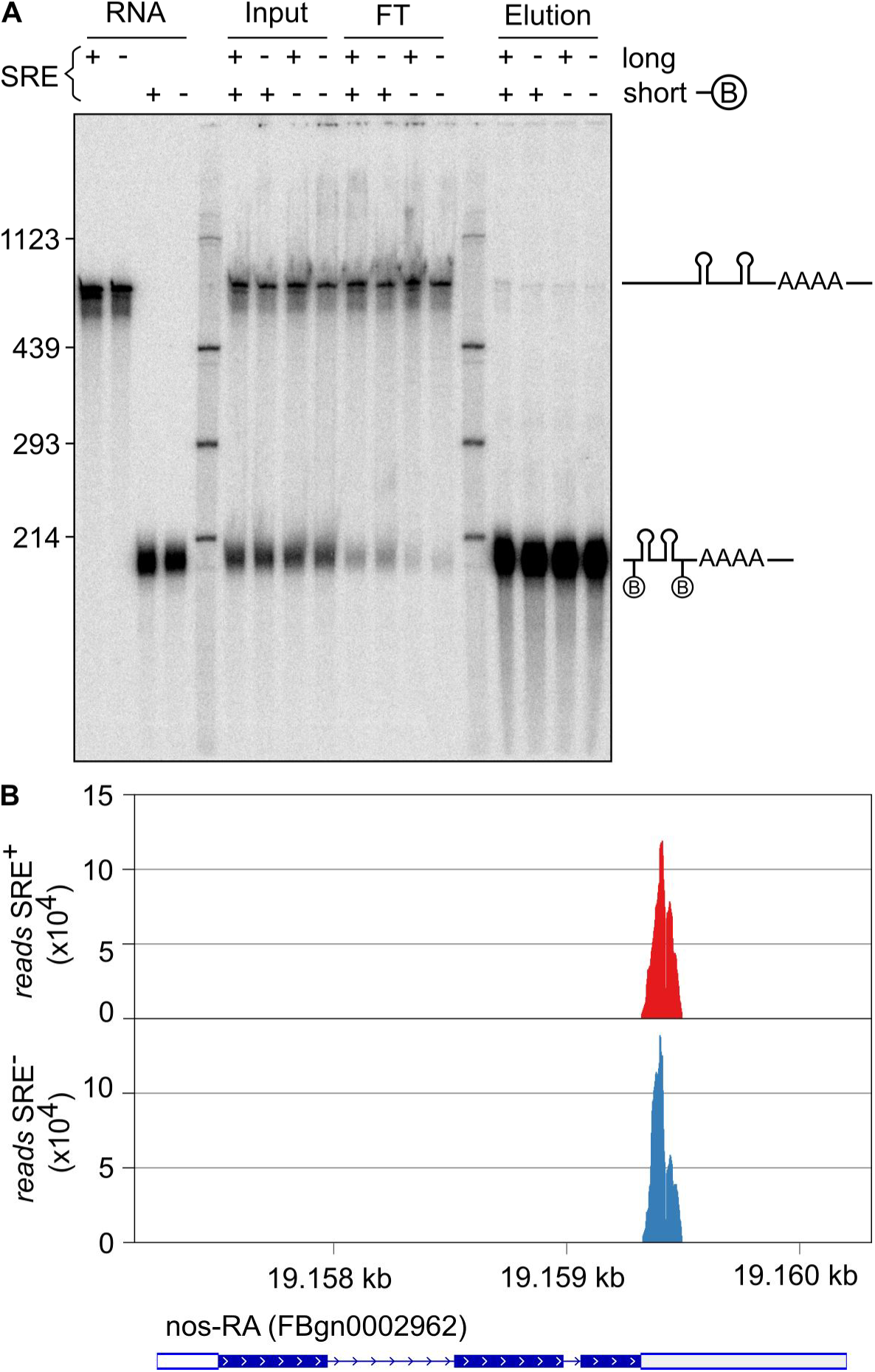
SRE-containing RNAs do not oligomerize. (A) A biotinylated RNA of 200 nt (SREonly; SRE^+^ or SRE^−^) and a non-biotinylated RNA of 630 nt (AUGonly; SRE^+^ or SRE^−^) were incubated together in embryo extract under conditions permitting assembly of the repressor complex. Streptavidin pull-downs were performed to enrich the biotinylated RNA together with potentially associated RNAs. RNA was eluted in formamide loading buffer at 95°C. The lanes labelled ‘RNA’ show the purified RNAs used, ‘input’ shows the RNAs after incubation in extract, ‘FT’ is the flow-through of the pull-down, and ‘elution’ shows the bound fraction. The figure shows one experiment of two. (B) RNA was purified from affinity-purified SRE^+^ and SRE^−^ RNPs and deep-sequenced. Reads mapping to the *nos* gene are displayed. For the experiment, bait RNAs were used at 10 nM. The abundance of *nos* has been estimated as 2 nM (Trcek et al. 2015). With a ~ twofold dilution upon extract preparation and an additional twofold dilution in the assay, endogenous *nos* sequences should have been detectable if an association with the bait RNA had taken place.

As an unbiased test for a potential association of the SRE^+^ RNA with other RNAs, total RNA was isolated from purified repressor complexes and from SRE^−^ controls and analyzed by deep sequencing. Although sequencing was targeted to small RNAs, *nos* sequences were also recovered; these were limited to the part contained in the bait RNA, and no difference between SRE^+^ RNA and SRE^−^ control was observed (**Fig. 4B**). Calculations (see figure legend) indicate that endogenous *nos* RNA present in the extract would have been detectable if it had been associated with the bait RNA. Thus, the repressed *nos* RNP does not oligomerize with other repressed RNPs. The lack of *nos* oligomerization is consistent with *in vivo* data (Little et al. 2015). No SRE-dependent enrichment of other RNAs was observed, making it unlikely that trans-acting RNAs are involved in SRE-dependent repression *in vitro*.

### Multiple copies of Me31B and Tral associate with the repressed RNA

Me31B orthologues can oligomerize on their own or when bound to RNA, and the ability of protein variants to oligomerize correlates with their ability to repress translation. The proteins appear to bind in multiple copies along RNA *in vivo* (Minshall and Standart 2004; Ernoult-Lange et al. 2012).

In order to determine whether oligomerization of repressor proteins on the reporter RNAs might play a role in SRE-dependent translational repression, we estimated the stoichiometries of proteins in the repressor complex: Three different biotinylated radiolabeled RNAs were used, each containing two copies of the SREs: SREonly (200 nt), the 1-AUG *nos* RNA used for the MS analysis (630 nt) and the luciferase reporter RNA (1956 nt); corresponding SRE^−^ RNAs served as controls. All RNAs were allowed to assemble repressor complexes before affinity purifications were carried out. The quantities of immobilized RNAs were determined from their specific radioactivities, and amounts of associated proteins were estimated by western blotting and comparison to standard curves of purified recombinant material. Representative data are shown in **Figs. 5A** and **S2**, and a summary of the average stoichiometries is presented in **Fig. 5B**. RNA association of Smg was SRE-dependent, but independent of RNA length: The stoichiometry was between 1 and 2 for all three RNAs. Within the accuracy of the experiment, this was equimolar with the SREs (see legend to **Fig. S2**). Binding of Cup was also approximately stoichiometric with the SREs. Bel bound independently of RNA length, but tended to be less abundant; with the longest RNA, specific binding was no longer distinguishable from background. In contrast to Smg, Cup and Bel, both Tral and Me31B clearly bound in a length-dependent manner, in excess of Smg and the SREs and approximately equimolar to each other. As these data indicate binding of multiple copies of Me31B and Tral along the RNA, it is unclear whether the amounts associated with the SRE^−^ RNAs should be subtracted as background or not. Without background subtraction, the stoichiometry of Me31B/Tral binding to RNA was near one copy of Me31B and Tral per 100 nucleotides. ‘Coating’ of RNA by Me31B and Tral may form an inert, ‘masked’ RNP that is at the core of translational repression, sterically preventing ribosome access to the RNA. For want of a reagent more comparable in size to a ribosome (3 × 10^6^ Da), accessibility of the repressed RNA was probed with the endonuclease RNase I (as an MPB fusion protein; 72,000 Da). The repressed SRE^+^ RNA proved to be considerably more resistant to the nuclease than the SRE^−^ control (**Fig. 5C, D**). This agrees with an earlier observation that an SRE^+^ RNA, when simply incubated in embryo extract under conditions of unchecked endogenous nuclease activity, was moderately more stable than an SRE^−^ control (Jeske et al. 2011). These data strongly argue in favour of sequestration of the RNA by a protein complex.

**Fig. 5:**
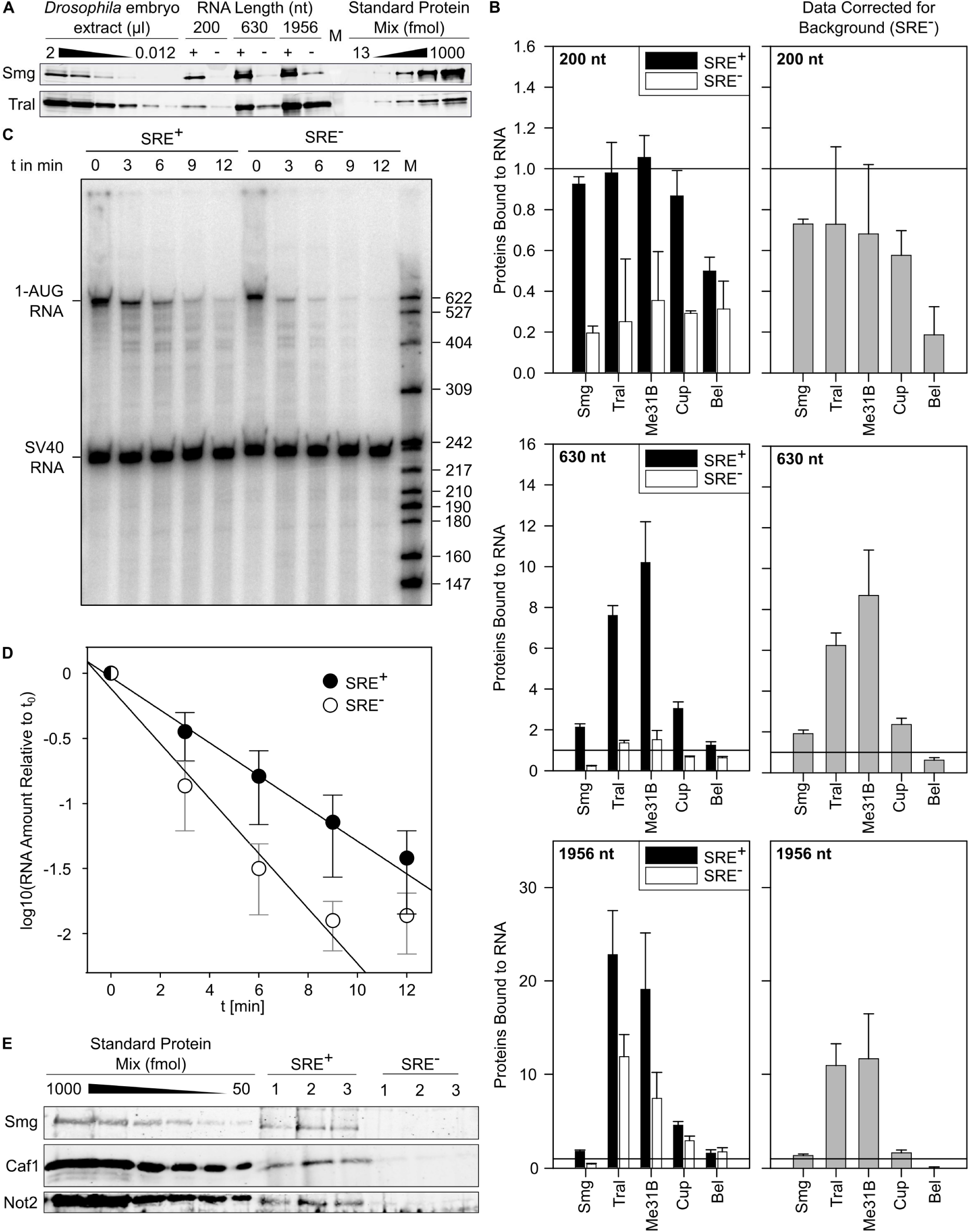
The SRE-dependent repressor complex sequesters the RNA through multiple copies of Me31B and Tral. (A) Three biotinylated RNAs of different lengths but each containing two SREs were used, together with matching SRE^−^ controls, for repressor complex formation in embryo extract and streptavidin pull-down. Bound proteins were analyzed by western blotting. Known amounts of recombinant proteins were used as standards. Analyses of Smg and Tral are shown as representative examples. (B) Stoichiometries of bound proteins were estimated from experiments as in (A). The panel on the left shows signals for SRE^+^ and mutant controls separately; in the panel on the right, protein binding to the SRE^−^ RNA was subtracted from the SRE^+^ signal. The horizontal lines mark a 1:1 molar ratio of protein to RNA. Error bars represent the standard deviation from 3–5 independent experiments. Additional data are presented in **Fig. S2**. (C) An RNase I protection experiment was carried out as described in Materials and Methods. (D) Quantification of experiments as shown in (**C**) (average of n = 4 with three independent batches of embryo extract). Error bars represent the standard deviation. Data were fitted to a first-order decay with the last time point of both RNAs omitted. The half-life of the SRE^+^ RNA was 1.7fold longer than that of the SRE^−^ control. (E) The association of Caf1 and Not2 with the SRE^+^ RNA and SRE^−^ control was examined as in (A). Three streptavidin pull-down experiments were carried out with the 630 nt RNA and independent batches of embryo extract. Western blotting and comparison to standard curves was carried out for the proteins indicated. The average amount of Smg recovered was 200 +/− 100 fmol. Tral was recovered at 1000 +/− 150 fmol in the SRE^+^ sample and at 500 +/− 120 fmol in the SRE^−^ sample (data not shown). All three subunits of the CCR4-NOT complex were present below the smallest amount in the standard curves (50 fmol). In a separate western blot, signals for Caf1, Not2 and Ccr4 were below 12.5 fmol (data not shown).

The stoichiometries indicate that Me31B and Tral cooperate as a defined subcomplex within the repressor complex. Indeed, treatment of a Smg immunoprecipitate with the cross-linker BuUrBu (Muller et al. 2010) identified a cross-link between Tral and Me31B consistent with the interaction dependent on the FDF motif of Tral (Tritschler et al. 2008; Tritschler et al. 2009) (**Figs. 6A, S3**). Thus, this interaction is likely to be relevant within the context of the repressor complex. When Tral and GST-Me31B were co-expressed in insect cells by means of baculovirus vectors, Tral was co-purified with GST-Me31B on glutathione beads, suggesting the existence of a stable complex (**Fig. 6B, C**). Co-purification was not affected by elevated salt concentration or RNase A.

**Fig. 6:**
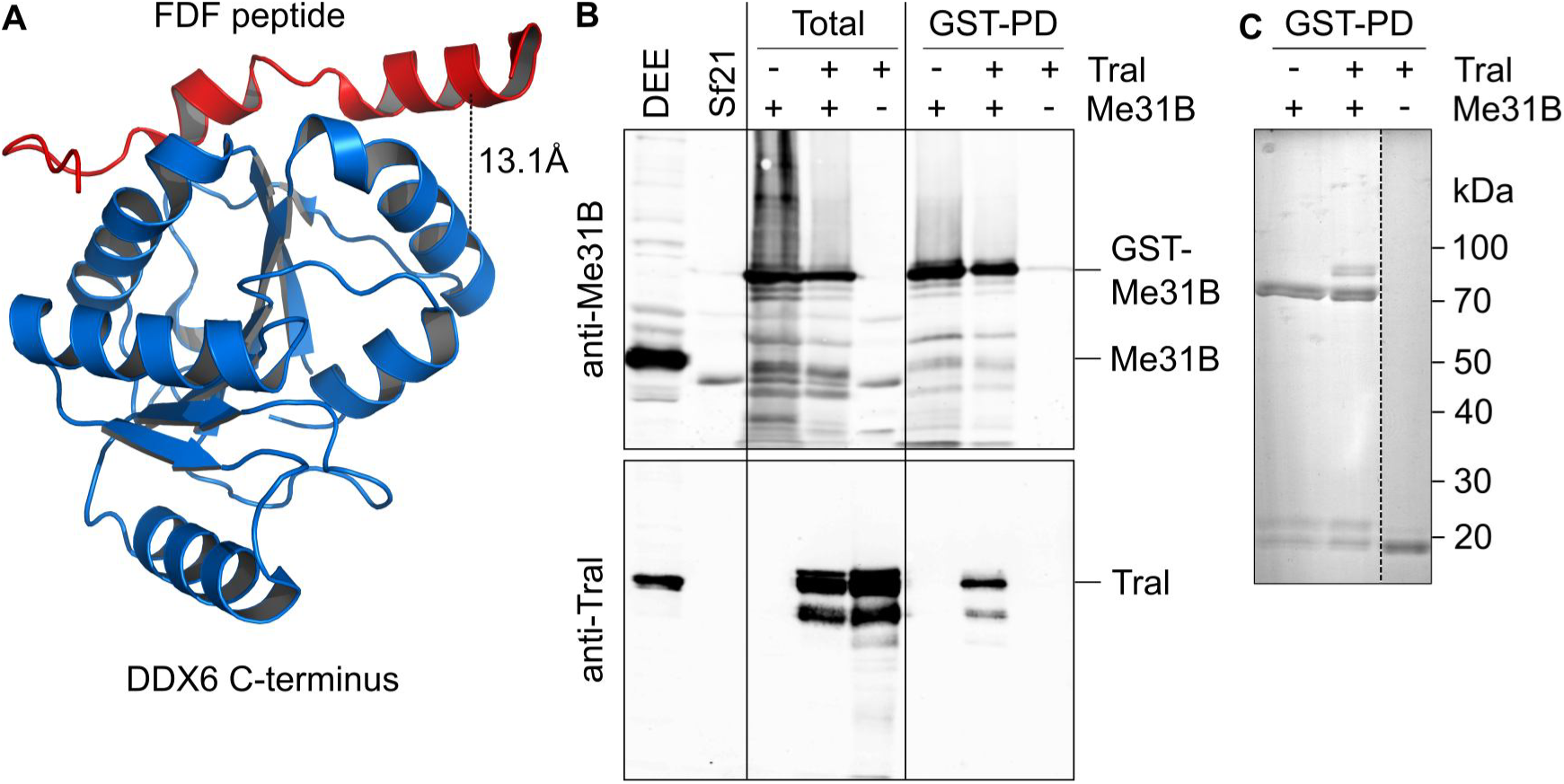
Me31B and Tral form a complex. (A) Structure of a complex between the C-terminal domain of DDX6 and an EDC3 peptide containing the FDF motif (PDB 2WAX) (Tritschler et al. 2009). Tral uses the same motif to bind Me31B. The black line represents the cross-link identified (**Fig. S3**) with the C_α_-C_α_ distance indicated. (B) Sf21 cells were infected with baculoviruses expressing GST-Me31B, Tral, or both as indicated. ‘Total’ refers to an SDS lysate. Purifications on glutathione beads were carried out from native lysates. Proteins were analyzed by western blotting for Me31B (top) or Tral (bottom). *Drosophila* embryo extract (DEE) and non-infected SF21 cells served as controls. (C) Glutathione bead eluates were analyzed by SDS polyacrylamide gel electrophoresis and Coomassie staining.

The components of the repressor complex were abundant among the soluble proteins of the embryo extract, as estimated by quantitative western blotting (with an error of ~ two; see Materials and Methods): Smg was present at 0.08 μM; Cup, 2 μM; Bel, 0.8 μM; Me31B, 3.5 μM, Tral, 7.6 μM. We estimate that extracts were ~ twofold diluted compared to egg content. In comparison, an mRNA concentration of roughly 0.4 μM in a *Drosophila* egg can be estimated on the basis of an egg volume of 0.01 μl (Azevedo et al. 1996), a total RNA content of 0.19 μg per egg (Hough-Evans et al. 1980), and the assumption that 2% of this is mRNA with an average length of 3000 nt. The ratio of protein to RNA concentration is consistent with Smg acting on a sizeable fraction of maternal mRNAs (Tadros et al. 2007; Chen et al. 2014a) and with Cup participating in translational repression exerted by other RNA binding proteins, e. g. Bruno (Nakamura et al. 2004; Wilhelm et al. 2005; Chekulaeva et al. 2006). The abundance of both Me31B and Tral is consistent with the two proteins binding in multiple copies and as a complex to repressed mRNAs. The high concentration of Me31B, exceeding that of mRNA, is consistent with data in other organisms (Ernoult-Lange et al. 2012) (and references cited therein).

The MS data suggest that the CCR4-NOT complex is not part of the core repressor complex. Association of the CCR4-NOT complex with the 630 nt RNA was examined by quantitative western blotting. In agreement with the MS analysis, CCR4, Caf1 and Not2 bound the RNA in an SRE-dependent manner, but were clearly substoichiometric (**Fig. 5E and data not shown**).

### Bel is required for *nos* mRNA translational repression *in vivo*

Belle is a DDX3-type RNA helicase. These proteins have been reported to be involved in translation, but both activating and repressive roles have been described. To examine a direct role of Bel in *nos* mRNA control in the embryo, we used two strong or null alleles of *bel, bel^6^* and *bel^L4740^*, which cause larval lethality. In addition, the hypomorphic allele *bel^neo30^* was used, which leads to female sterility when combined with stronger alleles (Johnstone et al. 2005; Ihry et al. 2012). Consistent with this, transheterozygous *bel^neo30/6^* and *bel^neo30/L4740^* females were sterile: When crossed with wild type males, they produced embryos (referred to as *bel^neo30/6^* and *bef^neo30/L4740^* embryos) that failed to eclose (**Fig. 7A**). *bel^neo30/6^* embryos showed a stronger phenotype than *bel^neo30/L4740^*, most of them being fragile and having short or no dorsal appendages.

**Fig. 7:**
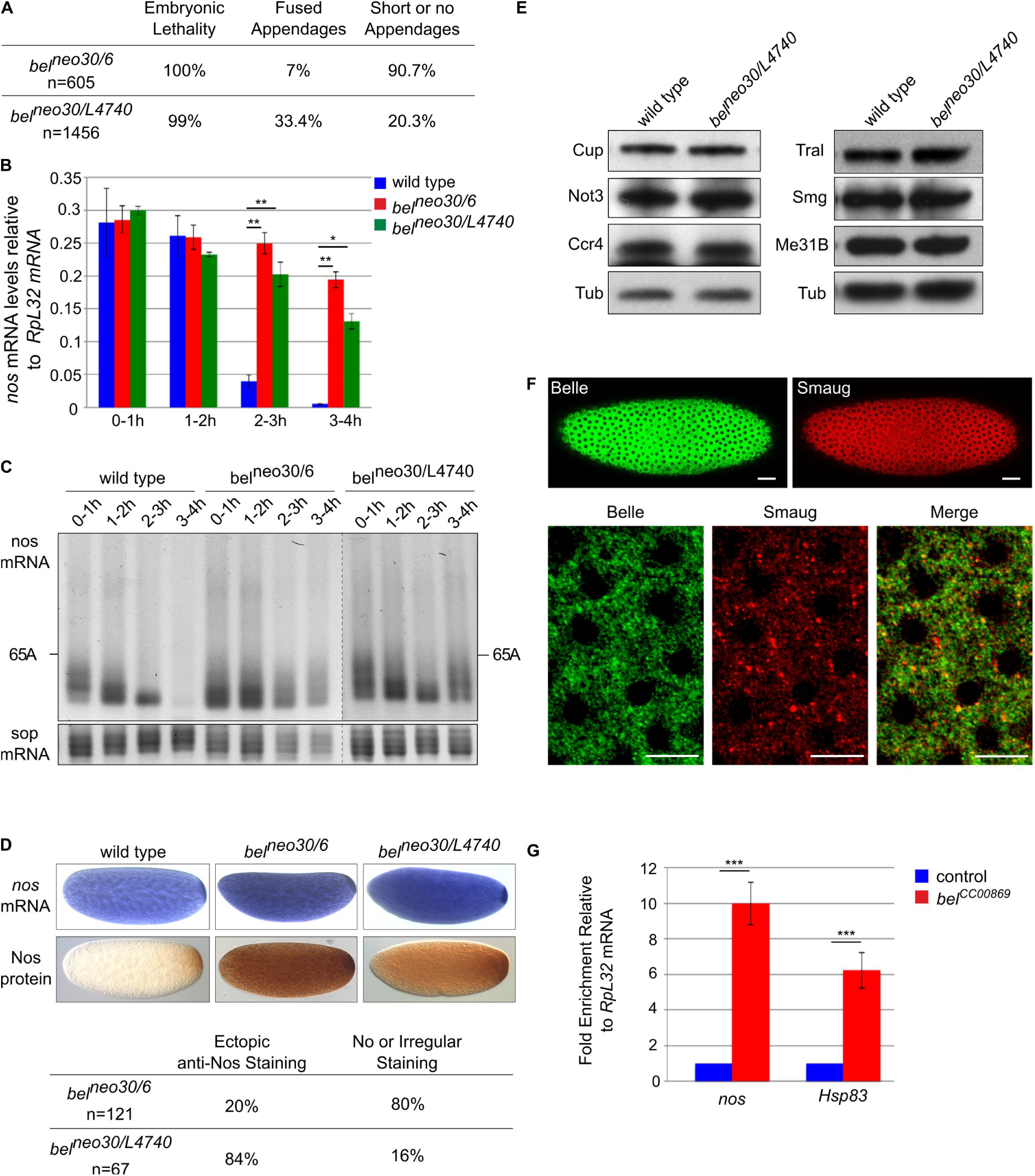
Bel is required for *nos* mRNA translational repression *in vivo*. (A) Phenotypic quantification of embryos coming from *bel^neo30/6^* or *bel^neo30/L4740^* mutant females crossed with wild type males. Numbers refer to the embryos examined. (B) *nos* mRNA quantification using RT-qPCR in wild type and *bel* mutant embryos spanning 1 h intervals up to 4 h of development. *RpL32* was used as a control mRNA for normalization. Means are from 3-4 biological replicates. The error bars represent SEM. * p<0.05; ** p<0.01 using the bilateral Student’s t-test. (C) PAT assays measuring *nos* mRNA poly(A) tail lengths in wild type and *bel* mutant embryos spanning 1 h intervals up to 4 h of development. PAT assay profiles using ImageJ are shown in **Fig. S4A**. *sop* encodes a ribosomal protein and was used as a control mRNA. (D) In situ hybridization of *nos* mRNA (top panels) and immunostaining with anti-Nos (bottom panels) of wild type and *bel* mutant embryos. Quantification of immunostaining is indicated below the images. (E) Western blots of wild type and *bel* mutant 0–2 h embryos probed with antibodies against six components of the *nos* repressor complex, including the CCR4-NOT complex. Anti-α-tubulin (Tub) was used as a loading control. (F) Confocal images of syncytial embryos co-stained with rabbit anti-Bel and guinea pig anti-Smg. Bottom panels show a higher magnification. Quantification using the Pearson Correlation Coefficient (PCC) indicated significant partial colocalization (PCC=0.52). Anterior is to the left. The scale bars represent 30 μm and 10 μm in top and bottom panels, respectively. (G) Quantification of *nos* and *Hsp83* mRNAs using RT-qPCR in anti-GFP immunoprecipitations from *bel^CC00869^* embryos that express GFP-Bel and control (wild type) embryos that do not express GFP. *RpL32* mRNA was used for normalization. mRNA levels in control embryos were set to 1. Means are from two biological replicates quantified in triplicates. The error bars represent SEM. *** p<0.001 using the bilateral Student’s t test.

To address a role of Bel in *nos* mRNA deadenylation and decay, we quantified *nos* mRNA by RT-qPCR in wild type and *bel* mutant embryos spanning 1 h intervals during the first 4 h of embryogenesis. *nos* mRNA decay was prominent after 2 h in wild type embryos, but was strongly impaired in *bel* mutant embryos (**Fig. 7B**). Accordingly, poly(A) test assays, used to measure *nos* mRNA poly(A) tail lengths in embryos up to 4 h of development, showed that deadenylation was inhibited in *bel* mutant embryos (**Figs. 7C, S4A**). *In situ* hybridization of 0–2 h embryos suggested that *nos* mRNA stabilization might even occur before 2 h of embryogenesis, since the staining was darker in *bel* mutants than in wild type embryos (**Fig. 7D**). Translational repression of nos was also impaired in *bel* mutant embryos: Immunostaining with anti-Nos antibody revealed ectopic, increased Nos levels in *bel^neo30/L4740^* embryos (**Fig. 7D**). *bel^neo30/6^* embryos showed heterogeneous staining: a large proportion (80%) were irregularly or not stained, but the remaining 20% showed again high levels of ectopic Nos protein throughout the embryo (**Fig. 7D**). The heterogeneity in *bel^neo30/6^* embryos could be due to earlier defects during oogenesis (Johnstone et al. 2005) or to a potential gain-of-function nature of the *bel^6^* allele: *bel^6^* has a stop codon after the first third of the coding sequence, which encodes a 4E-BP domain (Yarunin et al. 2011; Ihry et al. 2012). Thus, a truncated protein in *bel^6^* might dominantly affect translation through binding to eIF4E. Analysis of Nos protein levels by western blots were in agreement with the staining pattern of the majority of embryos, showing increased levels in 0–2 h *bel^neo30/L4740^* embryos and reduced levels in 0–2 h *bel^neo30/6^* mutant embryos (**Fig. S4B**). Defects in *nos* regulation in *bel* mutant embryos did not result from reduced levels of other components of the repressor complex or of the CCR4-NOT complex (**Fig. 7E**).

These results show that Bel participates in the repression of *nos* mRNA in the somatic part of the embryo and thus imply that Bel is present there. A GFP-tagged Bel protein has been reported to be distributed throughout the syncytial embryo (Johnstone et al. 2005). Immunostaining of embryos with anti-Bel and anti-Smg antibodies validated the cytoplasmic distribution of Bel throughout the embryo and its partial colocalization with Smg (**Fig. 7F**).

In an independent experiment to ask whether *nos* mRNA is bound to Bel in embryos, we used the GFP protein-trap *bel* allele *bel^CC00869^*, in which GFP is inserted in frame in the N-terminal part of Bel (Buszczak et al. 2007). RNA immunoprecipitation with anti-GFP antibody showed an enrichment of *nos* mRNA over *Rpl32* mRNA in 0–2 h embryos expressing GFP-Bel (*bel^CC00869^*) compared to control embryos. Another Smg target, *Hsp83* (Semotok et al. 2005), was also enriched (**Fig. 7G**).

Taken together, these results support a functional role of Bel in the *nos* repressor complex *in vivo*, acting on both translational repression and deadenylation.

## Discussion

We have identified seven stoichiometric components of the SRE-dependent repressor complex that are likely to explain its ATP-dependent formation, high stability and repressive potency: Smg, which directly recognizes the SREs; Cup, which associates with Smg; the DEAD-box ATPase Me31B and its partner Tral; a second DEAD-box ATPase, Bel; and finally the cap-binding initiation factor eIF4E and, with less certainty, the cytoplasmic poly(A) binding protein, PABPC. The repressor complex analyzed is functional since translation of the RNA on which it has assembled is fully repressed without further preincubation (**Fig. 1D**). Thus, assembly of the seven proteins identified constitutes the rate-limiting step of translation repression. The same complex likely facilitates deadenylation, since Smg and the SREs are also important for deadenylation of *nos* by CCR4-NOT. Accordingly, all core components of the CCR4-NOT complex were associated with the repressor complex.

The CCR4-NOT complex can also contribute to translational repression, independently of its deadenylase activity (Cooke et al. 2010; Braun et al. 2011; Chekulaeva et al. 2011; Kuzuoglu-Ozturk et al. 2016). However, CCR4-NOT was clearly substoichiometric and thus may not be essential for translational repression. The assay of the repressor complex only tests for constituents incorporated in the rate-limiting step, though; as the translation assay requires incubation of the gradient-purified repressed RNP with embryo extract, we cannot exclude that other components of the extract may associate with the stable complex and participate in its repressive activity, i. e. a protein that is not among the stable core components of the repressed RNP may still play a role in repression.

Five subunits of the conserved CTLH complex were enriched in the purified repressor complex, but substoichiometric with respect to the core components. The yeast edition of the complex is a ubiquitin ligase (Santt et al. 2008; Chen et al. 2017). Smg is degraded during cell cycle 14 (Dahanukar et al. 1999; Benoit et al. 2009), and most other core constituents of the repressor complex also strongly decrease during the maternal-to-zygotic transition (Gouw et al. 2009). The CTLH complex might be involved in the degradation of these proteins.

The DEAD box RNA helicase Bel was the only newly discovered constituent of the repressor complex. Enrichment of the protein was less pronounced compared to the other core components, but genetic data confirmed that Bel is required for both translational repression and deadenylation of *nos* mRNA *in vivo*. Bel orthologues Ded1p and DDX3 are known to be involved in translation, but their precise role is unclear, since both depletion and overexpression inhibit translation (reviewed in (Soto-Rifo and Ohlmann 2013; Sharma and Jankowsky 2014)). Bel and its orthologues can be localized in RNP granules containing repressed mRNAs and promote granule formation (Soto-Rifo and Ohlmann 2013; Sharma and Jankowsky 2014). Bel and *C. elegans* LAF-1 have been suggested to play a role in the translational repression of specific mRNAs, *bruno* and *tra-2*, respectively, but evidence for a direct role has been lacking so far (Goodwin et al. 1997; Yarunin et al. 2011). A cooperation of Ded1p with Dhh1p (Me31B) in translational repression is suggested by genetic and physical interactions (Tseng-Rogenski et al. 2003; Beckham et al. 2008; Drummond et al. 2011).

The idea that maternal mRNA in unfertilized eggs is not translated because it is masked by a ‘protective protein coat’ was proposed more than 50 years ago (Spirin 1966). Whereas mechanisms have been analysed that repress maternal mRNAs by targeting, directly or via the poly(A) tail, the 5’ cap function (Wilhelm and Smilbert 2005; Lasko 2011; Barckmann and Simonelig 2013), proteins coating and sequestering the RNA have not been identified with certainty. Circumstantial biochemical evidence has supported the concept, though: Repressed RNPs formed *in vitro* sediment rapidly, suggesting association of the RNA with many proteins and tight packaging (Chekulaeva et al. 2006) (this paper). Repressed RNAs are also moderately more stable in the face of nucleases endogenous to the extracts in which the assays were performed (Chekulaeva et al. 2006; Jeske et al. 2011). Repression of CRPV IRES-dependent translation, which is independent of all initiation factors, is consistent with exclusion of ribosomes (Jeske et al. 2011). Here we present evidence suggesting that the protective protein coat is formed by a complex of Me31B and Tral. The SREs nucleate the assembly of multiple copies of a Me31B•Tral complex on the RNA. The orthologue of Me31B is a component of stored *Xenopus* oocyte mRNPs (Weston and Sommerville 2006). The presence of Me31B and its partner Tral in the repressor complex is also consistent with previous reports of these two proteins interacting and causing translational repression (see Introduction). DDX6-type proteins bind RNA even in the absence of ATP (Dutta et al. 2011; Ernoult-Lange et al. 2012; Sharif et al. 2013). Tral presumably contributes directly to RNA coating, as it contains two types of potential RNA binding domains, an N-terminal Lsm domain and two RGG domains. Evidence for RNA binding by Tral orthologues has been published (Audhya et al. 2005; Tanaka et al. 2006). As the complex affords protection even against a relatively small endonuclease, we propose that it prevents translation by sterically excluding ribosomes, in agreement with the original idea of masking (Spirin 1966; Spirin 1994). Specificity of repression for the *nos* RNA depends on sequence-specific binding of Smg, but on the basis of biochemical similarities we suspect that other repressors may use similar mechanisms (Chekulaeva et al. 2006; Minshall et al. 2007).

A conceptual assembly of the repressor complex (**Fig. 8**) starts with Smg binding to the SREs. Smg binds Cup, which, in turn, associates with the Lsm domain of Tral (Tritschler et al. 2008; Igreja and Izaurralde 2011). Tral uses its FDF motif to bind Me31B (Tritschler et al. 2008; Tritschler et al. 2009; Igreja and Izaurralde 2011) (this paper), but Me31B can also directly interact with Cup (Nishimura et al. 2015; Ozgur et al. 2015). The mechanism of Me31B•Tral polymerization remains to be analyzed. Cup also brings in eIF4E (Wilhelm et al. 2003; Nakamura et al. 2004; Nelson et al. 2004; Zappavigna et al. 2004; Igreja and Izaurralde 2011; Kinkelin et al. 2012). Bel may join the complex via interactions with eIF4E (Sharma and Jankowsky 2014) or Me31B (Drummond et al. 2011). Candidates for recruiting the CCR4-NOT complex include Cup (Igreja and Izaurralde 2011; Kamenska et al. 2014), Me31B (Chen et al. 2014b; Mathys et al. 2014; Rouya et al. 2014; Ozgur et al. 2015; Waghray et al. 2015) and Smg (Semotok et al. 2005; Zaessinger et al. 2006). Me31B and Tral are both present in the embryo at very high concentrations, and micromolar concentrations of Me31B or Tral (Scd6p) can inhibit translation non-specifically *in vitro* (Coller and Parker 2005; Nissan et al. 2010). It will be interesting to find out how assembly of the stable Me31B•Tral oligomer is restricted to SRE-containing mRNAs.

**Fig. 8:**
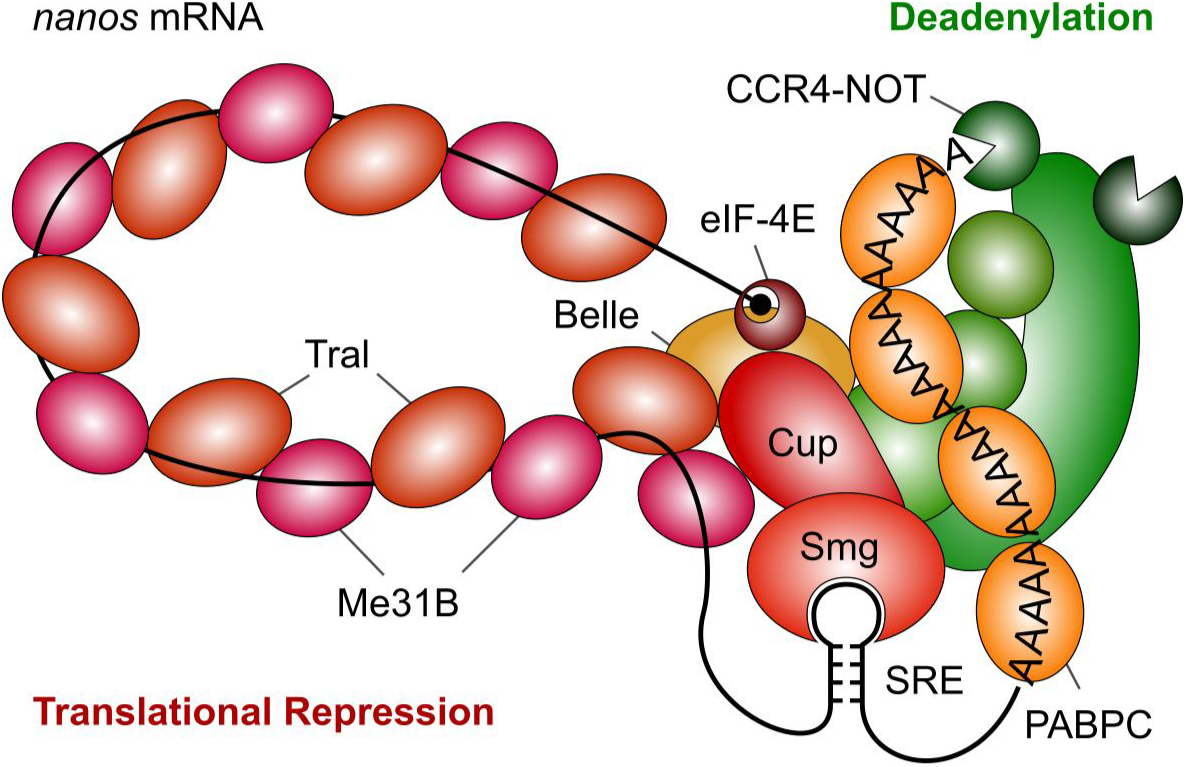
Model of the SRE-dependent repressor complex. The cartoon is based on the results of this paper and the references cited in the Discussion.

The presence of two ATP-dependent RNA helicases, Bel and Me31B, in the repressor complex probably accounts for its ATP-dependence. Me31B and/or Bel might also be responsible for the kinetic stability of the repressor complex. An attractive model is provided by the exon junction complex (EJC), which is frozen on the RNA because its central component, the DEAD-box ATPase eIF4AIII, is locked in a post-hydrolysis state by other EJC constituents (Ballut et al. 2005; Nielsen et al. 2009). Due to the cooperativity of ATP and RNA binding, this fixes the EJC on the RNA. Ded1 and two other DEAD box helicases tested were able to form very long-lived complexes with RNA in the presence of ATP analogs (Liu et al. 2014). Thus, this ‘clamping’ function may be a general feature of DEAD box helicases. We speculate that a component of the repressor complex may inhibit the dissociation of ATP or its hydrolysis products from Me31B and/or Bel to prevent the disintegration of the repressor complex.

Polymerization of Me31B and Tral along the RNA, nucleated by Smaug binding in the 3’ UTR, conceptually solves a problem that, to our knowledge, has barely been discussed in the literature, although it is faced by all 3’ UTR-bound protein complexes repressing translation initiation: Cartoons depicting the mechanism of action of such complexes invariably show an interaction of the 3’ end with the 5’ end, accompanied by the formation of an RNA loop. Any such interaction has to be intramolecular, i. e. the 3’ UTR-bound repressor complex has to find the 5’ end of its ‘own’ mRNA in the face of competition from ‘foreign’ 5’ ends. (For an interesting alternative, see (Macdonald et al. 2016).) One possibility for such an intramolecular interaction to occur would be ‘through space’: The two opposite ends of the flexible mRNA molecule diffuse randomly through the cytoplasm. As they are tethered to each other via the RNA body, an intramolecular interaction would be favored by a high local concentration of the *cis* 5’ end with respect to the regulatory 3’ UTR site. However, the efficiency with which this leads to an intramolecular interaction depends on variables like the length of the RNA and the concentration of competing 5’ ends. One would suspect that a more reliable mechanism should have evolved, in particular with a repressor complex as stable as the one described here: Any *trans* interaction established by mistake would not only result in a wrong RNA being repressed, but presumably also in a *nos* mRNA active in the wrong place. Polymerization of the Me31B•Tral complex along the RNA would constitute a fool-proof mechanism guaranteeing that the repressive action of the SREs is strictly intramolecular.

## Materials and Methods

### RNA

All RNA constructs (SRE only; 1-AUG *nos;* luciferase reporter; all with two wild type SREs or with a point mutation in each SRE) have been described (Jeske et al. 2006; Jeske et al. 2011). RNAs were synthesized with T3 RNA polymerase. Biotin-16-UTP (Jena Bioscience) and [α-^32^P]-UTP were incorporated during transcription at a reduced concentration of UTP. For incorporation of a similar number of biotin molecules per RNA, UTP was adjusted according to the number of uridines in the RNA (Luc: 1 mM, 1-AUG: 0.25 mM, SREonly: 0.1 mM) at a constant concentration of biotin-16-UTP (20 μM). When desired, an m7G cap was also incorporated co-transcriptionally. RNAs were gel-purified.

‘Short RNA’ was produced by partial hydrolysis of yeast RNA: 75 mg of yeast total RNA was dissolved in 5 ml 20 mM Tris-HCl, pH 8.0. 200 μl of 2.5 M NaOH was added, and the mixture incubated for 50 min at 40°C. 200 μl of 5 M HCl was added and incubation continued for 10 minutes at 40°C. After addition of 600 μl 3 M sodium acetate, the RNA was purified by phenol/chloroform extraction and isopropanol precipitation.

### Embryo extract and *in vitro* translation

Extracts were prepared as described (Jeske and Wahle 2008) except that embryos (Canton S) were 15 to 135 min old, and the lysate was centrifuged twice (20,000 x g, 30 min, 4°C). Aliquots were frozen in liquid N_2_ and stored at −80°C.

Luciferase reporter RNAs were incubated at 25°C in 40% embryo extract, 16 mM Hepes pH 7.4, 50 mM potassium acetate, 1 mM magnesium acetate, 0.8 mM ATP, 0.25 mg/ml yeast tRNA, 0.2 mg/ml ‘short RNA’, 0.08 g/l creatine kinase, 1 mM DTT, 80 U/ml RNAse inhibitor. Reactions were started with or without preincubation by the addition of 20 mM phosphocreatine and amino acids (20 μM each), incubated for 30 min at 25°C and stopped on ice. Luciferase activity was assayed with the Promega kit.

### Purification of the repressor complex

Radiolabelled, biotinylated RNAs (10 nM) were incubated under conditions in which no translation takes place (‘preincubation conditions’; 60% embryo extract, 26 mM Hepes-KOH pH 7.4, 81 mM potassium acetate, 1.6 mM magnesium acetate, 1.3 mM ATP, 1 mM DTT, 80 U/ml RNase inhibitor, 0.2 mg/ml ‘short RNA’) for 25 min at 25°C. The inclusion of ‘short RNA’ improved RNA stability and complex recovery. Aliquots (1 ml) were loaded on 5–45% sucrose gradients (12 ml per tube in TL buffer: 16 mM Hepes-KOH pH 7.4, 50 mM potassium acetate, 1 mM magnesium acetate, 0.8 mM ATP) and centrifuged for 3 h at 40,000 rpm, 4°C (Beckman SW40Ti). Gradients were harvested from the bottom in 20 fractions. Fractions 5-10 were pooled, frozen in liquid N_2_ and stored at -80°C. Pools from three gradients were combined and concentrated in Amicon centrifugal filters. Streptavidin beads (GE Healthcare; 30 μl packed volume) were blocked with TL buffer containing 0.1 mg/ml yeast RNA and 0.1 mg/ml methylated BSA and washed with TL buffer. Beads were incubated with equal amounts, based on trace-labeling of the RNA, of the concentrated pools and 0.1 mg/ml yeast RNA for 15 min at room temperature, pelleted and resuspended in 200 μl of TL buffer with yeast RNA as above. Beads were pelleted, resuspended in the same buffer and centrifuged through a 30% sucrose cushion in TL buffer (200 μl). They were washed once more with the same buffer in a fresh tube, once with wash buffer (50 mM Hepes-KOH pH 7.4, 150 mM KCl, 54 mM potassium acetate, 1 mM magnesium acetate, 30 μg/ml heparin, 0.1 mg/ml yeast RNA) and once with wash buffer without RNA. Proteins were eluted in 10 mM Tris-HCl pH 8.0, 0.5% SDS at 80°C for 10 min.

For ‘simple’ pull-down assays, the same procedure was used, but gradient centrifugation was omitted.

### Mass spectrometry

Streptavidin-purified repressor proteins from four preparations (twelve gradients) for each RNA were pooled and separated in an SDS-polyacrylamide gel. Each gel lane was cut in 12 pieces, and the proteins were in-gel digested with trypsin (Shevchenko et al. 2006). Disulfides were reduced with DTT and cysteines alkylated with iodoacetamide. Peptides were analyzed by LC/MS/MS on an U3000 RSLC Nano-HPLC system coupled to an Orbitrap Fusion Tribrid mass spectrometer equipped with a nano-electrospray ionization source (Thermo Fisher Scientific). The samples were loaded onto a trapping column (Acclaim PepMap C8, 300 μm x 5 mm, 5 μm, 100Å) and washed for 15 min with 0.1 % trifluoroacetic acid (TFA) at a flow rate of 30 μl/min. Trapped peptides were eluted on the separation column (Acclaim PepMap C18, 75 μm x 250 mm, 2μm, 100Å), which had been equilibrated with 99% A (0.1 % formic acid). Peptides were separated with a linear gradient: 0–35 % B (100% acetonitrile, 0.08 % formic acid) in 90 min at 40°C and a flow rate of 300 nl/min. Full MS spectra were acquired in the Orbitrap analyzer (R = 60,000), MS/MS spectra (HCD, 30% normalized collision energy) were recorded in the linear trap for 5 s (most intense signals).

MS data were analyzed with MaxQuant 1.5.2.8 (Cox and Mann 2008) (RRID: SCR:014485). For protein identification, data were searched against the Uniprot proteome (www.uniprot.org) of *D. melanogaster* (20,042 protein entries; accessed 2015/01/19). The inverted sequences of all proteins were used for decoy analysis. Mass accuracy was set to 20 ppm and 0.5 Da for precursor and fragment ions respectively. Carbamidomethylation was set as fixed modification and methionine oxidation and *N*-terminal acetylation were set as variable modifications. The search included standard contaminating proteins, but these were omitted from the plots shown. Raw files from the analysis of 12 gel pieces of one lane each for WT and MUT were combined into one experiment for MaxQuant analysis. The resulting peptide intensities for each protein were multiplied with the number of peptide spectral matches (PSMs) and normalized to the molecular weight of the protein. The calculated intensity values were plotted for proteins bound to SRE-containing RNA versus mutant RNA. For estimation of the p value for SRE-dependent enrichment, the proteins were separated in 33 equally sized bins along the diagonal axis. The distance of each protein from the diagonal in each bin follows a normal distribution with a mean of 0 for all nonspecifically bound proteins. The distances were fitted against the normal distribution to obtain σ^2^. To estimate p values for each protein enrichment, the σ^2^-values were fitted with the equation y=ab^-c(bin-45)^. The squared differences to the model were multiplied with the number of proteins per bin to weight the data in the non-linear least square fit.

### Analysis of RNA associated with the repressor complex

Repressor complex was isolated as above from 400 μl of reaction mixture without gradient centrifugation, and no yeast RNA was used during the pull-down and washing procedures. RNA was eluted with Trizol for 10 min at 80°C. 500 ng of total RNA was used in the small RNA protocol with the TruSeq™ Small RNA sample prepkit v2 (Illumina) according to the manufacturer’s instructions. The barcoded libraries were size restricted between 140 and 165bp, purified and quantified using the Library Quantification Kit - Illumina/Universal (KAPA Biosystems). Library pooling, cluster generation, high-throughput sequencing of 2x100bp and demultiplexing of raw reads was done according to (Stokowy et al. 2014).

Reads were stripped of the 3’ linker (TGGAATTCTCGGGTGCCAAGGAACTCCAGTCAC) using Cutadapt, and the resulting RNA sequences were mapped to the *Drosophila melanogaster* genome using Bowtie (100% match; release 5). Reads were first annotated to tRNA, rRNA, snoRNA, snRNA and miRNAs. piRNAs were the remaining reads that were 23–29 nt in length. piRNAs were mapped to TE using Bowtie with up to 3 mismatches. Uniquely mapped piRNAs were mapped to piRNA clusters using cluster coordinates from (Brennecke et al. 2007). mRNA-derived small RNAs were uniquely mapped reads that mapped in sense orientation to genes. Small RNA counts were normalized to 1 million mapped reads.

### RNase protection assay

2 nM radiolabelled 1-AUG-RNA (SRE^+^ or SRE^−^) was incubated under preincubation conditions (without DTT and RNase inhibitor) for 25 min at 25°C. 70 μL of the reaction was mixed with 35 μL of RNase I_f_ (NEB) at a final concentration of 0.66 U/μL. 15 μL aliqouts of the reaction were stopped at different time points in SDS-containing 2x proteinase K buffer with 20 μg Proteinase K, 20 μg glycogen and an unrelated radiolabelled RNA as extraction control. After incubation at 37°C for 30 min, the sample was ethanol precipitated and analysed on a denaturing 5% polyacrylamide gel.

### Western blots and immunostaining

The western blots in **Figs. 7 and S4** and immunostaining of embryos were performed as described (Benoit et al. 2005). In other experiments, SDS-polyacrylamide gels were blotted overnight in 25 mM Tris, 192 mM glycine onto PVDF-membranes and blocked in 5% milk in TBST. After primary antibody incubation, blots were washed with TBST and incubated with fluorescently labelled secondary antibodies (IR-Dye; LI-COR), washed and scanned on a LICOR scanner. The following proteins were used as standards for quantitative western blots: Me31B, Tral and Bel were the *E*. coli-produced proteins used for immunization (see below). His-tagged Ccr4, Not2 and Caf 1 were produced in E. coli from pET19 and purified under denaturing (Ccr4, Not2) or native conditions (Caf1). Flag-Smg and Flag-Cup were coexpressed in the baculovirus system and affinity-purified as a mixture. All standard proteins were quantitated by Coomassie staining of SDS-polyacrylamide gels and comparison to an unrelated standard protein; dye binding to different proteins under acidic conditions has been estimated to vary by about a factor of 2 (Chial and Splittgerber 1993). The Me31B standard was used to estimate the Me31B concentration in a batch of embryo extract which was then used as a standard in the analysis of pull-down assays. All other standard proteins were used directly.

Antibodies against Cup were obtained from Akira Nakamura (Nakamura et al. 2004) or Robin Wharton (Verrotti and Wharton 2000); against Nos from Akira Nakamura; against PABPC from Nahum Sonenberg (Imataka et al. 1998) or Matthias Hentze (Duncan et al. 2009); against Yps from James Wilhelm (Wilhelm et al. 2000). Antibodies against Me31B (Nakamura et al. 2001; Harnisch et al. 2016), Smg (Chartier et al. 2015), Ccr4, Caf1 (affinity-purified), Not2 (Temme et al. 2004) and Not3 (Jeske et al. 2006) have been described. Additional guinea pig antibody against Smg was from Craig Smibert (Tadros et al. 2007). Antibodies against Bel were initially obtained from Paul Lasko (Johnstone et al. 2005), and antibodies against Tral were initially from Elisa Izaurralde (Tritschler et al. 2008). Additional antibodies against Bel and Tral were generated as follows: N-terminally His-tagged variants of the proteins were expressed in *E. coli*. Cells from a 400 ml culture were resuspended in 100 mM Tris-HCl pH 7.0, 1 mM EDTA, incubated with 1.5 mg lysozyme, and lysed by ultrasonification. After DNase I treatment of the lysate, 20 mM EDTA, 2% Triton and 500 mM NaCl were added, insoluble proteins were pelleted for 10 min at 31,000 g and washed with 100 mM Tris-HCl pH 7.0, 20 mM EDTA. The pellets were dissolved in urea buffer (8 M urea, 100 mM Tris-HCl pH 8.0, 100 mM Na_2_HPO_4_) and proteins bound to Ni-NTA-Agarose. The columns were washed with urea buffer and urea buffer with 20 mM imidazole and proteins eluted with urea buffer plus 250 mM imidazole. After concentration with AMICON centrifugal filters, the proteins were diluted with PBS to less than 4 M urea, and 800 μg was used by Eurogentec S.A. (Belgium) to immunize two rats (Tral) or two rabbits (Bel).

For MS analysis of Smg-associated proteins, 2 x 60 μl of Protein A-Sepharose beads (GE Healthcare) were washed 3 times with TL-buffer (see: Purification of the repressor complex). Beads (60 μl) were incubated with 20 μl of anti-Smaug serum or the corresponding preimmune serum in 500 μl TL-buffer for 2 h at ~8°C, washed twice with wash buffer (see: Purification of the repressor complex) and twice with TL-buffer. The beads were incubated with 500 μl *Drosophila* embryo extract, diluted 1:4 in TL-buffer for 2 h at ~8°C, transferred to Protein-LoBind tubes (Eppendorf) and washed with TL-buffer (2 times), wash buffer (3 times) and TL-buffer (2 times). In one sample, bound proteins were cross-linked for 2 h at 8°C with 0.5 μM BuUrBu cross-linker (Muller et al. 2010) in 500 μL of TL-Buffer. Bound proteins were denatured in 100 μl 8 M urea, 0.4 M ammonium bicarbonate, disulfides were reduced with 10 mM DTT and alkylated with 55 mM iodoacetamide. The sample was diluted to 0.8 M urea and digested with trypsin overnight at 37°C. LC/MS/MS analysis was performed as described above except that MS/MS spectra (HCD, 30% normalized collision energy) were acquired in the Orbitrap analyzer (R = 60,000) for 5 s (most intense signals). Data were evaluated as described above. Cross-links were analyzed by means of the MeroX software (Gotze et al. 2015) with the following parameters: Trypsin was set as the protease, the BuUrBu cross-linker was selected, methionine oxidation was considered as variable modification, cysteine alkylation was set as static modification, the signal-to-noise ratio was set to 1.5, precursor precision was set to 3 ppm and fragment ion precision was set to 20 ppm. The analysis was performed with activated RISE-mode, and data were searched against a sequence database containing the set of enriched proteins from the RNA-pull-down experiment as well as sequences of 200 standard protein contaminants (cRAP-database http://www.thegpm.org/crap/index.html), which resulted in one highly confident cross-linked peptide pair between Me31B and Tral.

For RNA immunoprecipitations, 50 μl Protein A Mag Sepharose beads (GE Healthcare) were pre-washed twice in RIP buffer (25 mM Hepes pH 6.8, 250 mM sucrose, 1 mM MgCl2, 1 mM DTT, 150 mM NaCl, 0.1% Triton-X100, containing freshly added complete protease inhibitor cocktail EDTA free (Roche) and RNasin (Promega)). 10 μl of mouse anti-GFP (monoclonal antibody 3E6; Invitrogen) were added, the mixture was incubated in 500 μl RIP buffer for 2 h at 4°C on a wheel, and beads were washed twice in RIP buffer. Embryos (0–2 h old) were homogenized on ice in four volumes of RIP buffer and incubated for 20 min on ice. The homogenate was centrifuged twice at 10,000 g for 5 min and pre-cleared on 50 μl of RIP buffer-washed Protein A Mag Sepharose beads for 45 min at 4°C on a wheel. Beads were removed, and the extract was mixed with the anti-GFP antibody beads and incubated for 1.5 to 2.5 h at 4°C with rotation. The beads were washed eight times with RIP buffer, extracted with Trizol, and the RNA was isopropanol-precipitated in the presence of glycogen and resuspended in 12 μl H_2_O.

### Expression of GST-Me31B and Tral

The coding sequence of Me31B with an N-terminal GST tag and PreScission site was cloned into pFastBac1 and transferred into the Bac-to-Bac baculovirus expression system (ThermoFisher). Expression clones for untagged Tral were generated by the same procedure. For protein expression, Sf21 cells were infected at MOI = 1 and harvested three days later. For western blots, equal numbers of cells were lysed in SDS gel loading buffer. For GST pulldown, cells were sonicated in GST buffer (50 mM HEPES pH 7.5, 150 mM KCl, 1.5 mM MgCl_2_, 10% saccharose). The cleared lysate was incubated with Glutathione Sepharose (GE Healthcare) for 2 h at 4 °C. After incubation, beads were washed three times briefly and two times for 10 min in GST buffer. Bound proteins were eluted in SDS gel loading buffer and analysed by western blot or SDS-PAGE and Coomassie staining. Copurification of Tral with GST-Me31B was not affected by the inclusion of RNase A or elevated salt concentration (400 mM KCl).

### Fly stocks

The *w^1118^* stock was used as control. *bel* mutant alleles were *bel^6^*, *bel^L4740^* and *bel^neo30^* (Bloomington Stock Center). The GFP protein-trap allele *bel^CC00869^* corresponds to a GFP insertion after the first coding exon at amino acid 15; it contains the complete Bel coding sequence (Buszczak et al. 2007).

### *In situ* hybridization, PAT assays and RT-qPCR

Whole-mount in situ hybridization was performed by standard methods. The probe was an antisense RNA made from the pN5 *nos* cDNA clone (Wang and Lehmann 1991). Poly(A) test (PAT) assays and RT–qPCR were performed as described (Rouget et al. 2010) on two to four independent RNA preparations. Reverse transcription for qPCR was done using random hexamers (Invitrogen) and Superscript-III (Invitrogen). Real-time PCR (qPCR) was performed with the LightCycler System (Roche Molecular Biochemical) using *RpL32* as a control mRNA. Primers were as follows: Hsp83-fw-qPCR: CAACAAGCAGCGTCTGAAAAG; Hsp83-rev-qPCR: AGCCTGGAATGCAAAGGTC; nos1128F: CGGAGCTTCCAATTCCAGTAAC; nos1281R: AGTTATCTCGCACTGAGTGGCT; RpL32F: CTTCATCCGCCACCAGTC; RpL32R: CGACGCACTCTGTTGTCG; nosPostPAT: TTTTGTTTACCATTGATCAATTTTTC; sopPAT: GGATTGCTACACCTCGGCCCGT

### Microscopy and Image Processing

Fluorescent images were acquired using a Carl Zeiss LSM 780 LASER scanning confocal microscope (Montpellier RIO Imaging facility) and a 40X PLAN Apochromatic 1.3 oil-immersion objective. The acquisition software was Zen. Contrast and relative intensities were processed with ImageJ software. Light microscope images were acquired using Leica Leitz DMRB Fluorescence Microscope with Nomarsky lens. Colocalization was quantified by FIJI as follows: background was substracted with a rolling ball radius of 30 μm, and the Pearson Correlation Coefficient (PCC) was calculated using the colloc2 plugin with auto-thresholding. Mean PCC was calculated from three images.

## Acknowledgments

We thank F. Kluge and D. Taenzler for excellent assistance with experiments; P. Enke for generating baculovirus clones; K. Krohn, IZKF, University of Leipzig, for performing Illumina sequencing; M. Hentze, E. Izaurralde, P. Lasko, A. Nakamura, C. Smibert, N. Sonenberg, R. Wharton and J. Wilhelm for reagents; H. Ashe for fly stocks; F. Falvo for the BuUrBu crosslinker; and M. Jeske and C. Eckmann for comments on earlier versions of the manuscript. This work was supported by grants from the DFG (WA 548/13–2 within FOR 855; WA 548/16–1; GRK 1591) to EW and by CNRS-University of Montpellier UMR9002, ANR (ANR-15-CE12-0019-01), FRM (Equipe FRM 2013 DEQ20130326534) and Fondation ARC (ARC Libre 2009, N°3192) to MS. JD was supported by Fondation ARC.

